# Lipid Droplet metabolism dependent microbial defense in pre-immune zebrafish embryos

**DOI:** 10.1101/478859

**Authors:** Asmita Dutta, Sampali Banerjee, Deepak Kumar Sinha

## Abstract

Microbes present survival challenge to pre-immune embryos. Our study provides evidence for antimicrobial-secretion-based strategy of zebrafish embryos against microbes during pre-immune stages. Chorion prevents physical contact between embryos and microbes, yet microbes compromise embryonic survival through their secretions. Development of embryos in microbe-free medium involves secretion of pro-microbial compounds that bacteria utilize to accelarete growth. Embryo senses presence of microbes through microbial secretions. They respond by altering their secretions to include antimicrobial compounds along with regular pro-microbial ones. Upon sensing embryonic anti-microbial secretions, microbes too alter their secretions to include more potent toxins for embryos. In response to this embryos alter their secretions to include more potent antimicrobial compounds. Ability of embryos to secrete antimicrobial compounds is positively correlated with amount of lipid droplets (LDs) in them. Inhibition of LD metabolism prevents antimicrobial secretions by embryos. Thus, LDs protect zebrafish embryos from microbes. This manuscript establishes that pre-immune embryos employ dynamically evolving biochemical warfare to protect themselves from harmful microbes.

## Introduction

The immune systems of fish and mammals share significant similarities. For example, both the vertebrates host comparable sets of lymphocytes (similar T-cell subsets) and hence, generate analogous immune responses^1–3^. However, unlike mammalian embryos that develop inside the body of the female, embryonic development of fish (oviparous) occurs in the open aquatic environment. Therefore, right after fertilization these oviparous embryos require a functional immune system to safeguard them from pathogenic microorganisms present in their natural habitat. Studies on zebrafish embryos report that adaptive immune response develops four to six weeks after fertilization of the egg^4–6^. Further, 'primitive macrophages' appear only after 16 hours post fertilization (hpf)^7–9^. These macrophages and the newly generated neutrophils constitute the innate immune system in one day old zebrafish embryos^2,10^. Hence, these embryos are exposed to a large number of pathogens prior to the development of either the innate or the adaptive immune system. However, it is not known how the zebrafish embryos protect themselves from pathogens before 1 day post fertilization (dpf) when even the innate immune system is not functional. Lipid droplets (LDs) are fat reservoirs with a neutral lipid core delimited by a phospholipid monolayer studded with various proteins^11–13^. Recent investigations have recognized the importance of LDs beyond lipid metabolism and homeostasis. They exhibit other non-canonical functions^14–16^ in addition to coordination of immune responses^17^. The lipid and the protein content of the LDs determine the inflammatory activity of the myeloid cells, thereby suggesting a crucial role in determining the cellular functionalities^18,19^. LDs in *Drosophila* embryos are reported to host anti-bacterial properties. In addition to LDs, the antimicrobial phosvitin (Pv), a nutritional protein abundant in eggs, protects the embryos from microbes.^20^ In zebrafish embryos, the chorion prevents all physical contact between the embryo body and the microbes thus the antimicrobial properties of LDs or Pv are not useful in dealing with the extra-embryonic microbes. The pathogenic microbes in the aqueous environment however challenge the embryos by releasing secretions which are permeable to the chorion. Strategies by the embryos to counter pathogenic challenges remain unknown. In this manuscript we explore how the embryos oppose the pathogens during early development (i.e. until 9 hpf). Our results indicate that LD metabolism is central to the embryonic strategies (until 9 hpf) that mitigate challenges posed by pathogenic microbes in aquatic environment.

## Results

### Mother zebrafish alone dictates the embryonic LD density (LDD)

The zebrafish embryos contain large number of LDs distributed cortically in the blastodisc (Fig. 1A). Fig. 1B compares the density of LDs (the number of LDs/mm^2^ of the blastodisc) in the embryos obtained from three different sets of parents. In accordance to our previous report^21^, the LDD increases with the development of the embryos, however, we also observe a unique LDD_30_ value (LDD at 30mpf) associated with each parent set (Fig. 1B, Supp. S1). Interestingly, the rate-of-increase of LDD (slope) with embryonic development, for embryos from a particular parent set, exhibits a strong positive correlation (correlation coefficient=0.75) with the corresponding LDD_30_ values (Fig. 1C). Therefore, we propose that the rate of synthesis of the LDs depends upon the initial LDD values (LDD_30_), which in turn depends upon the respective parent set used for breeding. Next, we explored whether the unique embryonic LDD_30_ is determined independently or jointly by the male and the female parents. We found that the LDD_30_ of the embryos obtained from independent females bred with different males do not alter LDD_30_ significantly (Fig. 1D), while similar analysis with the same male fish bred with different females show significantly varying LDD_30_ values in the offspring (Fig. 1E). Hence, embryonic LDD is uniquely determined by the female parent alone. This allowed us to obtain the embryos with distinct LDD by selecting the appropriate females for breeding.

**Figure 1.**
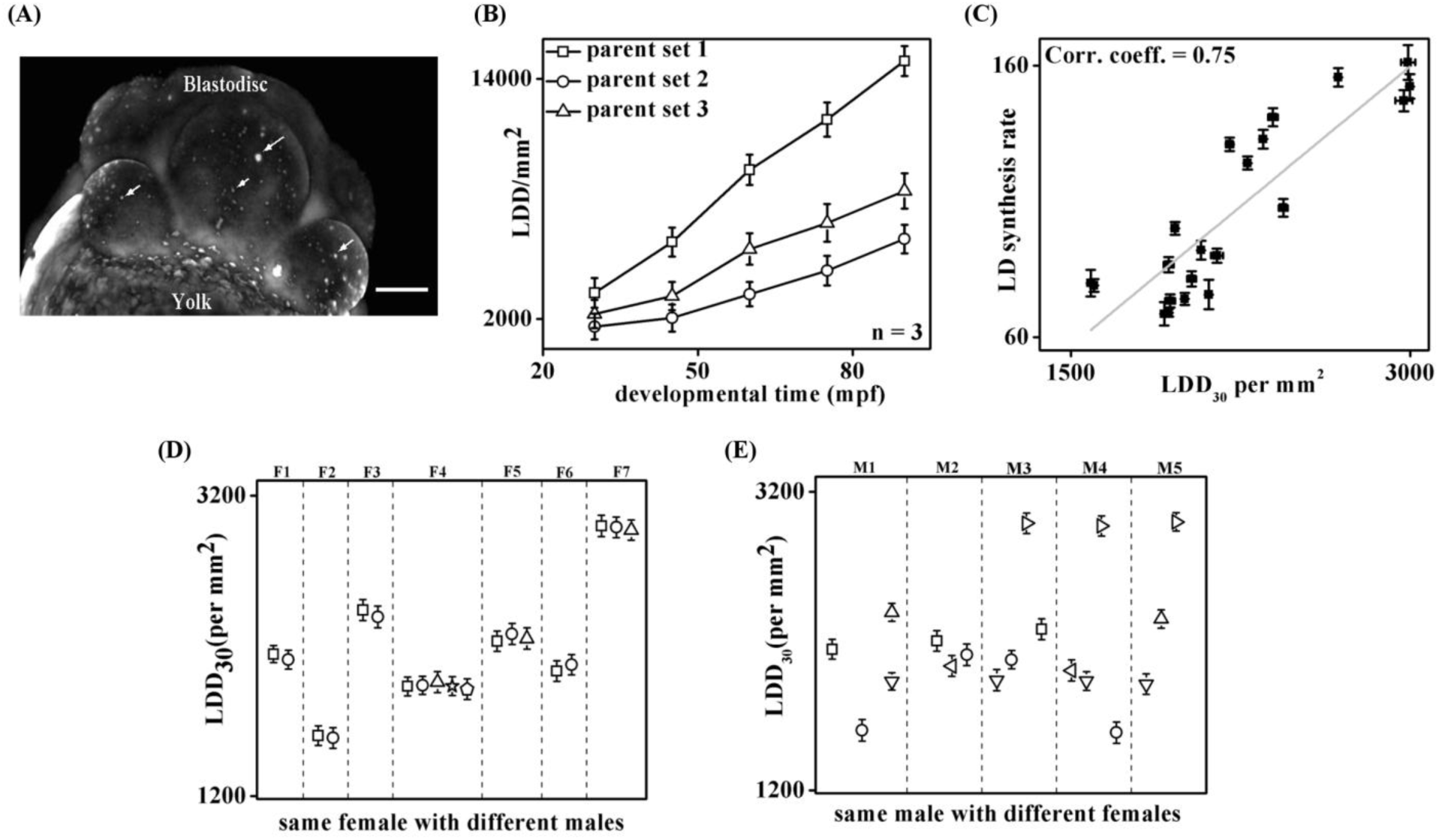
Embryonic LDD is a maternal characteristic. **(A)** Fluorescence image of 8-cell stage embryo stained with LD540, arrows show LDs, scale bar 100 μm. **(B)** Average LDD for a minimum of five embryos each, from three representative parent sets plotted against developmental time (mpf). Error bars are obtained from embryos acquired across different clutches from the same parents (n=3). **(C)** LD synthesis rate (rate-of-increase of LDD) plotted as a function of respective embryonic LDD_30_ values. Fit line denotes a positive correlation (0.75). **(D)** LDD_30_ re-plotted for 7 independent female parents (F1 to F7) bred with different males (different symbols). **(E)** LDD_30_ re-plotted for 5 independent male parents (M1 to M5) bred with different females (different symbols), error bars denote Standard Error of Mean (S.E.M.).

### LDs help the embryos endure microbial challenges

In the natural habitats, the zebrafish embryos are exposed to different microbes prevalent in the aqueous environment. These microbes do challenge survival of the embryos. Life expectancy (e_x_) of embryos is remarkably lower in water obtained from local pond (natural habitat) compared to sterile E3 (media prepared in laboratory) (Fig. 2A). To understand the impact of microbes on the survival of embryos, we incubated the embryos in bacteria-laced E3 media. For embryos with a given LDD, presence of bacteria (gram negative/gram positive) in E3 reduces their e_x_ in a concentration dependent manner (Supp. S2). We selected a bacterial concentration of 10^6^/ml at which the survival chances of embryos (LDD=2971) diminishes to 50% at 24 hpf (Supp. S2). We observe a similar reduction in likelihood of survival of embryos in E3 laced with bacteria (Fig. 2B). Interestingly, the embryos exhibit LDD_30_ dependent life expectancy in the presence of microbes (Supp. S3), as confirmed by the corresponding higher correlation coefficient (r=0.75 in presence of microbes) in Fig. 2C. Therefore, we hypothesize that the LDs are components of the antimicrobial defense system of the nascent zebrafish embryos. This hypothesis is further validated by injecting additional LDs into newly laid embryos and incubating them with/without bacteria. Exogenous LDs in the embryos increases their survival in bacteria-laced medium (Fig. 2D). However, exogenous LDs have no impact on the embryonic survival in bacteria-free medium (i.e. in E3) (Supp. S4). Hence, the LDs assist the embryos to counter the pathogenic challenge presented by the microbes.

**Figure 2.**
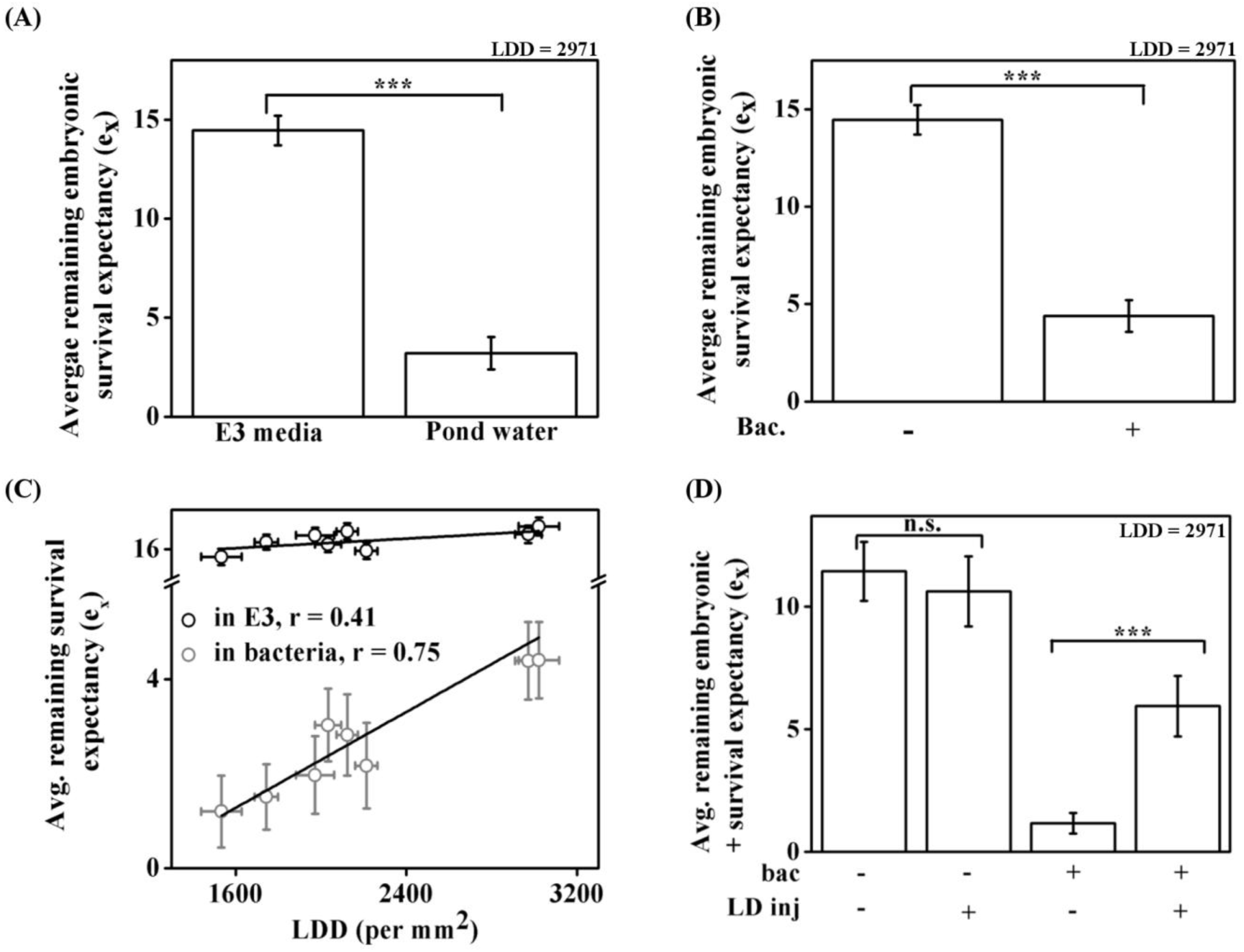
Survival of embryos in presence of microbes is dependent upon LDD. **(A)** Average remaining embryonic survival expectancy (e_x_) of embryos incubated in E3 and pond water (as mentioned in x-axis). **(B)** e_x_ of embryos incubated in the presence (left bar) or absence of bacteria (right bar) (as menitoned in x-axis). **(C)** e_x_ of embyros kept in E3 containing bacteria (gray) or in E3 alone (black) plotted against corresponding LDD_30_ values. Correlation fit shows a Pearson coefficient of 0.75 (with bacteria in E3) and 0.48 (in E3). **(D)** e_x_ of embryos incubated with/without bacteria in E3, with/without exogenous LD injection. LDD_30_ of the embryo clutch is 2971/mm^2^ in A, B, D. Bacterial concentration used: 10^6^/ml. All embryos incubated with external food supply so that bacteria is the only challenge to embryos.

### Lipolysis of LDs is necessary to confer defense against microbes

LDs present in the *Drosophila* embryos are reported to exhibit antimicrobial properties^22^. Our studies too reveal that the LDs isolated from zebrafish embryos host anti-bacterial property *in vitro* (Supp. S5). However, if the embryos utilize the antimicrobial property of the LDs to counter the microbial challenges then they must come in physical contact with the microbes. To test this possibility, we investigated if the bacteria in E3 have the ability to penetrate the chorion and the outer membrane of the embryos. For this the embryos were incubated in RFP-*E.coli* laced E3. Fig. 3A (i) establishes that the bacteria (in green) fail to penetrate even the outer chorion of the embryo (in gray). Though, the chorion prevents any physical contact between the embryo and the bacteria, yet presence of bacteria in E3 is deleterious to embryonic survival. This therefore, suggests that bacteria release permeable deleterious secretions that cause decrease in embryonic life expectancy. Since the chorion is the first line of defense that comes in physical contact with the microbes, we studied its antimicrobial activity. We find that though the chorion lacks cytotoxic capability, it is cytostatic (Fig. 3A (ii)). However, the antimicrobial capabilities of the chorion are not sufficient to protect the embryos from higher concentration of microbes. Next we investigated the effect of deleterious bacterial secretions (BS-E3: obtained by incubating bacteria in E3 overnight and filtering the supernatant; refer Table 1) on embryonic life expectancy (e_x_) as a function of embryonic developmental stage.

**Table 1.** Nomenclature for different media obtained from secretions under different conditions.

| Nomenclature | Secretion from | Incubation medium |
| --- | --- | --- |
| ES-E3 | Embryo | Embryonic secretions in E3 buffer |
| BS-E3 | Bacteria | E3 buffer |
| ES-BS | Embryo | (BS-E3): Bacterial secretions in E3 |
| ES-BS<sub>ES</sub> | Embryo | (BS-ES): Bacteria secretions in ES-E3 |
| ES-BS(OS) | Embryo | (BS-E3 + 10 μM OS): Bacterial secretions in E3 with OS |
| BS-ES | Bacteria | ES-E3 Embryonic secretions in E3 |
| BS-ES<sub>BS</sub> | Bacteria | ES-BS: Embryonic secretions in BS-E3 |

**Figure 3.**
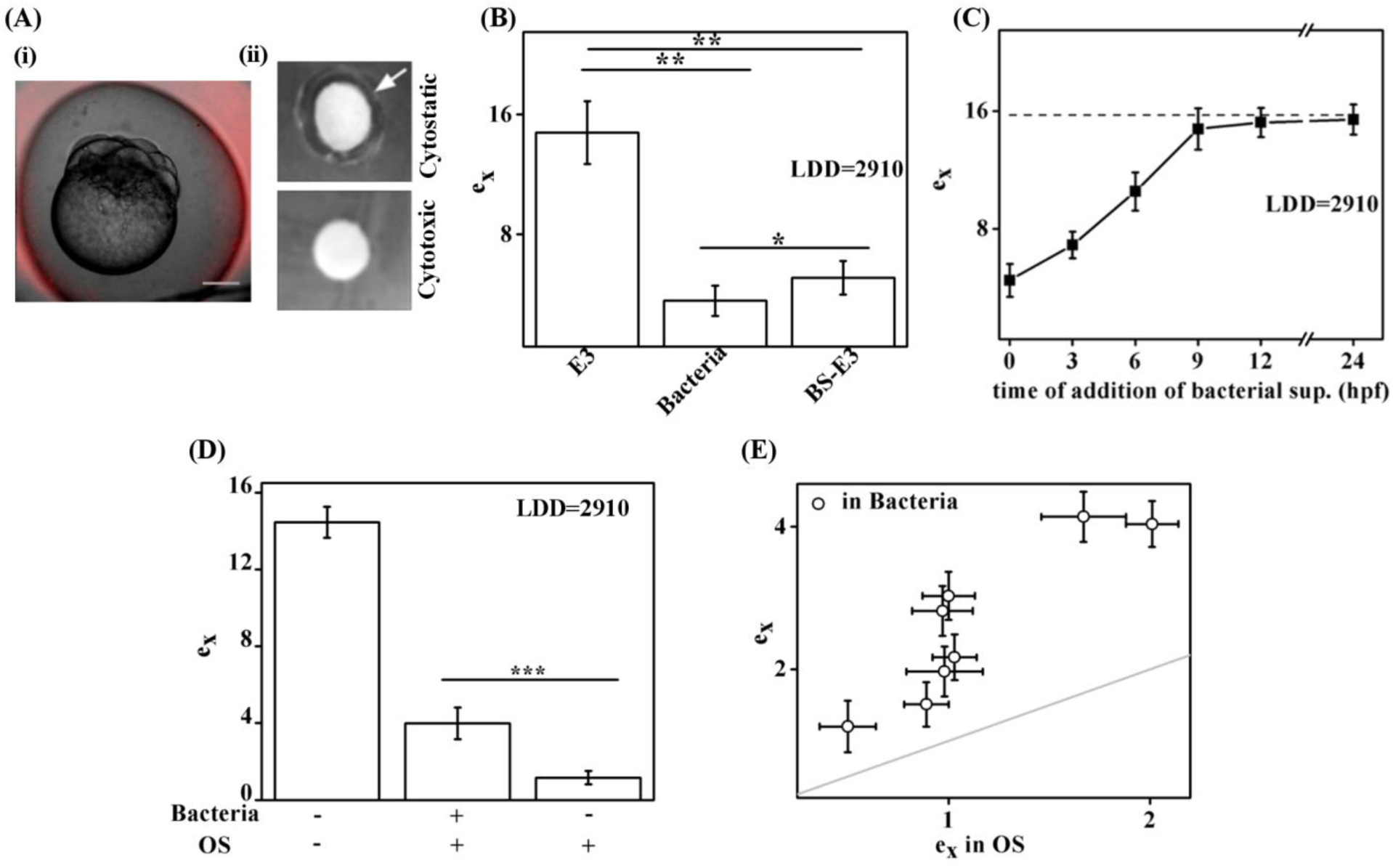
Embryonic survival under bacterial exposure is dependent upon LD-lipolysis. **(A) (A) (i)** Over-laid image of embyro (DIC) and RFP-*E.coli* (fluorescence) showing the inability of the bacteria to penetrate the chorion, scale bar 300 μm. **(ii)** Disc diffusion assay for cytostatic (upper) and cytotoxic (lower) property assesment of chorion against *E.coli*. Arrow: zone of inhibition. **(B)** e_x_ for embryos incubated in E3 (left bar), with bacteria in E3 (middle bar), in BS-E3 (right bar). **(C)** e_x_ for embryos incubated in bacterial supernatant added at different durations (x-axis denotes different time points when BS-E3 was added to embryos, in terms of hpf). Dash line denotes e_x_ of control embryos (grown in E3). **(D)** Plot of e_x_ verus the different conditons of incubation. The embryos were incubated in the presence/absence of bacteria and +/- OS in the medium. The embryo clutch used has LDD 2910 /mm^2^. **(E)** e_x_ of embryos plotted versus e_x_ in OS containing E3 for embryos incubated in bacteria (black). Different data point indicate embryos with different LDD_30_ values. Gray line denotes straight line (y=x).

Similar to incubation in bacteria-laced-E3, embryonic incubation in BS-E3 (a solution free of bacteria but contains bacterial secretions) also leads to significant decrease in the survival of the embryos (Fig. 3B). Embryos exposed to BS-E3 right from 0 hpf (1 cell stage) exhibit lower life expectancy as compared to the embryos exposed to BS-E3 at later developmental stages (i.e 9 hpf onwards, Fig. 3C). Since higher LDD bestows embryos with better capabilities to counter microbial challenge during 0-9 hpf (Fig. 2C), hence we investigated if metabolism of the LDs is required for such defense capabilities against microbes. The breakdown of LDs by the cytosolic lipases is one of the primary pathways for the catabolism of lipids and subsequent energy generation^12^. To investigate the role of LD metabolism in the survival of the embryos under bacterial invasion, we used the broad spectrum lipolysis inhibitor, Orlistat (OS)^23,24^, to reduce the activity of the embryonic lipases. Treatment of the embryos with OS at the working concentration severely retards the development of the zebrafish embryos^25^. Therefore, we titrated the concentration of OS to determine the appropriate concentration (10 μM) that does not affect the rate of development of the embryos significantly (Supp. S6). Irrespective of the LDD, inhibition of LD lipolysis by OS reduces embryonic life expectancy in bacteria-laced-E3 (Fig. 3D, 3E). Thus, metabolism of LDs is an integral part of the embryonic strategy to combat deleterious bacterial secretions, and mere presence of LDs is not sufficient to bestow embryos with microbe countering capabilities.

### Embryonic antimicrobial secretions mitigate the microbes challenge

Fig. 4A depicts the level of secretion obtained from embryos with distinct LDD_30_ values kept under different incubation conditions. We find that the level of embryonic secretions in E3 (ES-E3) is independent of LDD (r=0.089). However, incubation of embryos in BS-E3 leads to increase in the level of secretions for all the embryos (with different LDD_30_ values from different female parents) (Fig. 4A). As expected, additional embryonic secretion (ES-BS) is dependent on lipolysis of the LDs and inhibition of lipolysis by OS reduces the levels of embryonic secretions in BS-E3 incubation media (Fig. 4A). Thus, the embryos release lipolysis dependent additional secretions to counter the challenges presented by BS-E3. We had observed earlier that the survival of the embryos in bacteria is LDD dependent (Fig. 3C). This is consistent with the observation that the embryos release higher levels of secretions in BS-E3 if the embryonic secretions host antimicrobial property. Fig. 4B depicts the antimicrobial activity of embryonic secretions under different conditions. While ES-E3 is neither cytostatic nor cytotoxic (Fig. 4B-i), ES-BS does exhibit antimicrobial (cytostatic) activity (Fig. 4B-ii). As expected, ES-BS obtained from OS containing media does not host any anti-bacterial property (Fig. 4B-iii). Therefore, the embryos release LD-lipolysis dependent cytostatic secretions to mitigate the bacterial challenges in their surroundings. Since zebrafish embryos are susceptible to BS-E3 just until 9hpf (Fig. 3C), we investigated the cumulative embryonic secretions i.e. ES-BS until a certain developmental stage (or time in 9 hpf). Fig. 4C depicts the cumulative amount of secretions released by the embryos when exposed to bacterial secretions i.e. ES-BS until a given developmental stage. We observe that the embryos in E3 secrete mainly during the initial 3 hpf. There is no significant secretion 3hpf onwards (Fig. 4C). On the other hand, the embryos exposed to BS-E3 continue to release secretions until 9 hpf at a much higher rate. Interestingly, beyond 9 hpf neither are the embryos vulnerable to BS-E3 (Fig. 3C), nor do they secrete antimicrobial compounds (Fig. 4C). Next, we characterized the embryonic secretions using Thin Layer Chromatography (TLC), which can resolve the components present in the secretions based on their polarity. Fig. 4Di is a representative TLC profile when ES-E3 is run on a TLC plate and stained with Ninhydrin (for amino acid detection). The corresponding values of the resolving fractions (R_*f*_) for the spots were plotted in Fig. 4Dii. R_*f*_ defines the migration of a particular component on the TLC plate w.r.t. the migration of the solvent front. Similarly, the TLC profiles of the embryonic secretions under different conditions (ES-BS, ES-BS_ES_) (Table 1) have been represented in terms of their corresponding R_*f*_ values (Fig. 4E). The secretions by the embryos (ES-BS) incubated in BS-E3 (Fig. 4E, left panel of scatter plots) has significantly different TLC profile compared to ES-E3 (Fig. 4Dii). This suggests that embryos secrete many new compounds (e1-e7) in response to bacterial toxins (BS-E3). Most interestingly, the TLC profile of embryonic secretions, ES-BS, differ from that of ES-BS_ES_, (Fig. 4E, Suppl. S7), probably due to the different material compositions of BS-E3 and BS-ES. Thus, bacteria too have different secretions in E3 and ES-E3. As expected inhibition of LD lipolysis (by OS treatment) inhibits secretion of all compounds (e1-e7), instead we find an additional secretion spot (i.e. e8) that was not seen earlier. TLC plate of same samples in a non-polar solvent exhibit no spot with lipid-charring solution, thereby proving that the embryonic secretions are amine group containing compounds (probably amino acids) and not lipid based compounds. Both the secretions ES-BS and ES-BS_ES_ host cytostatic property (Fig. 4Fi, ii). However, inhibition of lipolysis results in the loss of antibacterial property of the embryonic secretions (Fig. 4Fiii). Release of amino acid based embryonic secretions upon BS-E3 exposure may be a result of either enhanced protein translation or active protein degradation, both of which are characteristic phenomena of early embryonic development. We determined an appropriate concentration of the protein translation inhibitor, Cycloheximide (CHX), at which the embryos survive however, they do show a reduction in the level of total protein due to CHX (1 μM) (Supp. S8). 1 μM of CHX does not result in any significant difference in the level of ES-E3 or ES-BS (Fig. 4G). Similarly, neither the treatment of Heclin (inhibitor of protein degradation) produced any significant difference in the level of ES-E3 or ES-BS (Supp. S9). Surprisingly, neither protein translation nor protein degradation seems to be the mechanism of synthesis of the cytostatic secretions (e1-e7). Next we explored the mechanism by which the microbes challenge the embryos.

**Figure 4.**
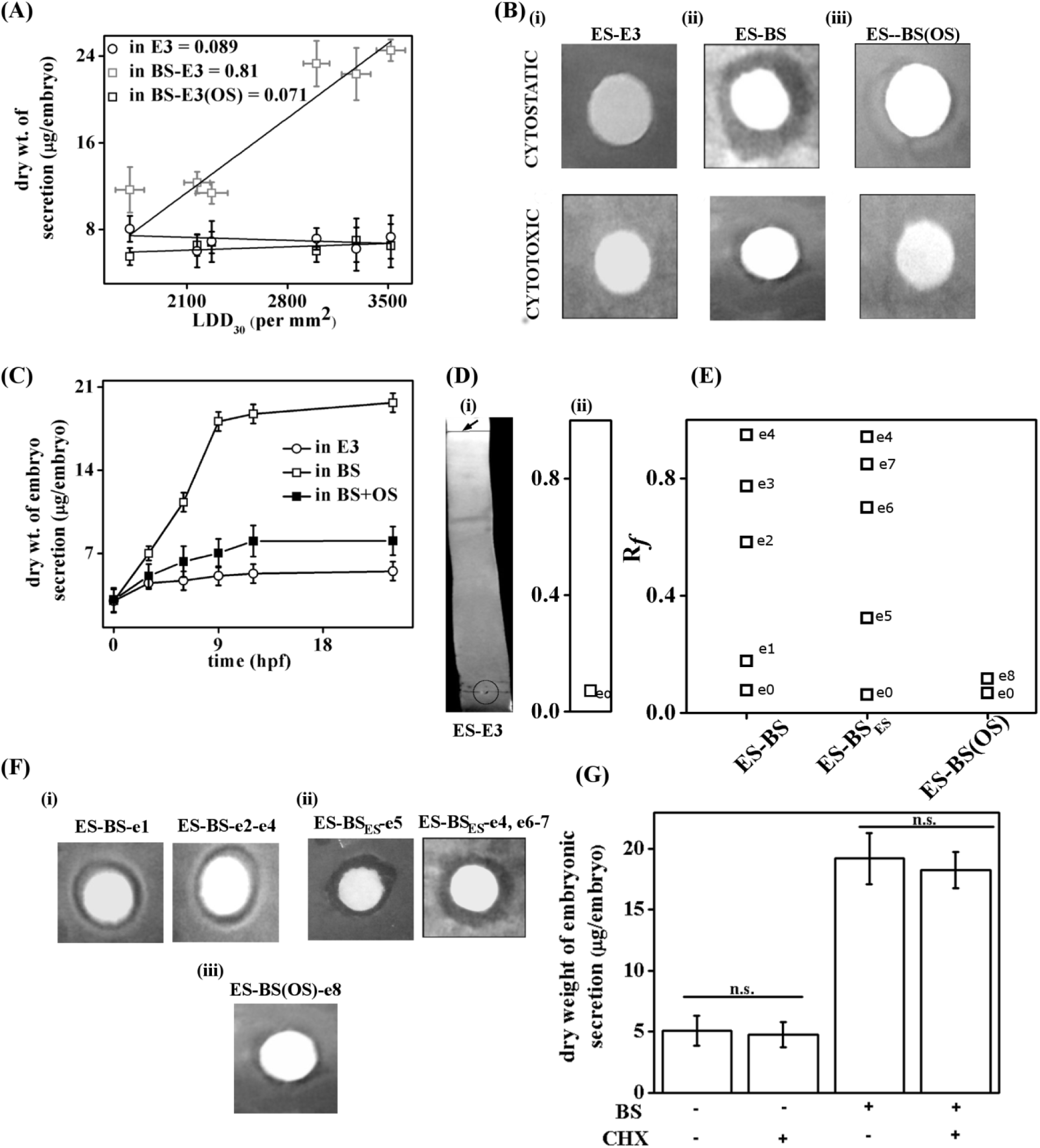
Lipolysis dependent secretion is released by embryos when incubated in BS-E3. **(A)** Dry weight of secretion (μg/embryo) of components when embryos are incubated in E3 (circle), BS-E3 (gray, square) and OS (10μM) added to BS-E3 (square, black) plotted against respective LDD_30_, correlation coefficient provided as legend. **(B)** Disc diffusion assay: cytostatic (upper panel) and cytotoxic (lower panel) property assesment of (i) ES-E3, (ii) ES-BS, (iii) ES-BS(OS). **(C)** Cumulative dry weight of embryonic secretion of components (μg/embryo) incubated in E3 (circle), BS-E3 (open, square) and OS (10 μM) containing BS-E3 (closed, square). Error bars denote S.E.M. **(D) (i)** TLC profile of ES-E3 stained by Ninhydrin. Arrow shows the solvent front, **(ii)** Corresponding R_*f*_ values for each spot on TLC plate plotted against the respective embryo incubation conditons. **(E)** Plot of R_*f*_ of TLC spots stained with Ninhydrin for embryonic secretions obtained from different incubation conditions as mentioned in x-axis. **(F)** Cytostatic property assessment of TLC-spots from (i) ES-BS, (ii) ES-BS_ES_ and (iii) ES-BS(OS) (numbering as per E). **(G)** Dry weight of secretion (μg/embryo) for embryos incubated with/without BS-E3 supplememted with/without CHX.

### Bacteria secretes more potent toxins in response to the embryonic secretions, ES-E3

We investigated the nature of the bacterial secretions under different incubation conditions. As expected, we observe a concentration dependent effect of BS-E3 on life expectancy (e_x_) of the embryos (Fig. 5A). The secretion, BS-E3, is obtained from bacteria that was allowed to grow in sterile E3. We compare the bacterial secretions in E3 (i.e. BS-E3) and in ES-E3 (i.e. BS-ES) by measuring their dry weight (Fig. 5B). Next we estimated the life expectancy of the embryos incubated in same amount of secretions released by bacteria under different incubation conditions. Fig. 5C confirms that BS-E3 (bacterial secretions in E3) and BS-ES (bacterial secretions in E3 with e0, refer Fig. 4D) have significantly different effects on the embryonic life expectancy. The bacteria secretes more potent toxins (BS-ES) upon exposure to ES-E3. The comparison of TLC profile of bacterial secretions under different conditions (column1 and coulmn2) (Fig. 5D) reveals that the bacteria secretes additional compound ‘b1’ only if they are exposed to ES-E3 (e0). More interestingly, bacteria exposed to compounds e0-e4 (antimicrobial secretions from embryos, refer Fig 4E) alter the composition of their secretions even further as the secretions in BS-ES are different from that in BS-ES_BS_ (Fig. 5D, b2-b4 are the additional bacterial secretion compounds, Suppp. S10). Additionally, these secretions are even more potent toxins for the embryos (Supp. S11). Like embryonic secretions, the bacterial secretions are too amine containing compounds and give rise to distinct spots in Ninhydrin stained-TLC plates. Next we investigated the effect of embryonic secretions on the bacteria. Fig. 5E depicts the growth curve for bacteria (*E.coli*) exposed to compound e0 (10^6^/ml bacteria grown in 2 ml ES-E3) obtained from embryos having distinct LDD_30_ values. Significantly, higher growth rate of bacteria in ES-E3 compared to E3 indicates that the compound e0 is a nutrient for the bacteria. Thus, normal development of zebrafish embryos involves secretion of e0 which aggravates the microbial challenge for the embryo itself by accelarating the bacterial growth rate. Fig. 5F depicts the growth curve for bacteria (*E.coli*) exposed to e0-e4 (10^6^/ml bacterial grown in 2 ml ES-BS) obtained from embryos having distinct LDD. We observe an inverse relation between the bacterial growth rate and the LDD of the embryos from which the ES-BS was obtained. This observation is in agreement with Fig. 4, since the amount of antimicrobial secretion (Fig. 4B) is positively correlated with LDD_30_ (Fig. 4A). We further validate this hypothesis by studying the bacterial growth rate in ES-BS (OS). Because of lack of antimicrobial activity of ES-BS (OS), the growth rate is no more dependent on LDD (Fig. 5G).

**Figure 5.**
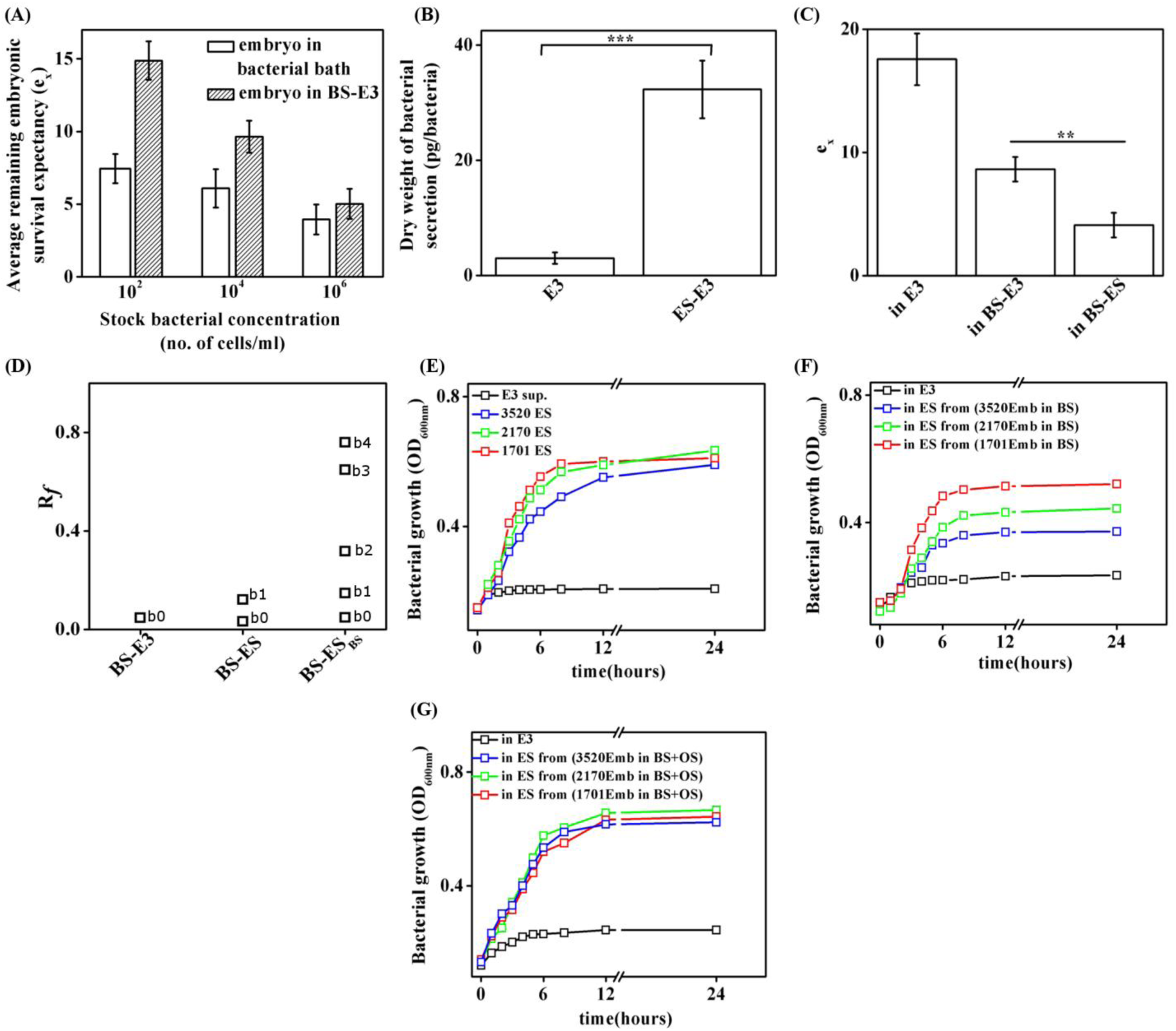
Mechanism of protection of embryos from bacteria by LDs. **(A)** e_x_ plotted against concentration of the bacterial stock. Embryos were incubated in either bacterial bath (empty bar) or in BS-E3 (filled bar). **(B)** Dry weight of secretion (pg/bacteria) by bacteria grown in E3 and ES-E3 plotted against the incubation conditions as mentioned in the x-axis. **(C)** e_x_ plotted against different incubation conditions as mentioned in the x-axis. Same amount of secretion (1 mg/ml) was suspended in E3 and then used to treat the embryos. **(D)** Plot of R_*f*_ of spots obtained by running TLC using BS-E3, BS-ES, BS-ES_BS_. Bacterial growth curves (O.D. at 600 nm) plotted against time (hours) for bacteria grown in **(E)** ES-E3 obtained from different embryo sets (with different LDD_30_), **(F)** ES-BS from different embryo sets, **(G)** ES-BS (OS) obtained from different embryo sets. Graph legends denote the LDD_30_ values of the embryo clutches.

## Discussion

Oviparous animals such as zebrafish fertilize their eggs externally. The fertilized eggs (embryos) face two types of survival challenges presented by their environment: 1) starvation and 2) pathogenic microbes. Relatively late onset of adaptive immunity in zebrafish embryos suggests their vulnerability during 0-9 hpf. Therefore, we investigated: (a) if the embryos are vulnerable to common microbes such as *E.coli*, and if yes, then (b) how do the embryos respond to pathogenic challenges during 0-9 hpf in the absence of any speciallized immune cells? Our observation that every clutch of embryos obtained from a given female fish has unique density of LDs (LDD) in them (Fig. 1), allows us to explore the link between LDs and survival of the embryos challenged with starvation or pathogen. LDs are synthesized on the Endoplasmic Reticulum (ER) surface. We speculate that the positive correlation in Fig. 1C arises probably from uniquely inherited ER content by the embryos. Therefore, both the LDD_30_ (amount of LD at 30 mpf) and the rate of LD synthesis are expected to be proportional to the ER surface area. By mimicking the pathogenic challenges present in the aquatic habitat (using the most ubiquitous microbe *E.coli*) we also show that the LDs help the embryos survive under bacteiral exposure (Fig. 2). The link between the abundunce of LDs and embryonic ability to cope with the microbes is surprising. Antimicrobial chorion of the zebrafish embryo (Fig. 3A-ii) which is the first line of defense against microbes, prevents any physical contact with microbes (Fig. 3A). The microbes attack the embryos through permeable toxic secretion (Fig. 5A). The embryos are susceptible to microbial secretions (b0, Fig. 5A) only until 9 hpf. The development of zebrafish embryos involves secretion of compound e0 (Fig. 4D) which is nutrient for bacterial growth (Fig. 5F). The embryos respond to bacterial toxin ‘b0’ by secreting antimicrobial amine containing compunds ‘e1-e4’ (Fig. 4E). Bacteria too sense the presence of embryos through embryonic secretion e0, and respond by secreting more potent pathogenic secretion ‘b1’ (Fig. 5B and D). Interstingly, ‘e0’ accelarates the bacterial growth rate (Fig. 5F). As a result more pathogenic secretion ‘b0’, ‘b1’ are generated, which compromise the survival of embryos even further. To counter the detrimental effects of pathogenic secretions ‘b0’, ‘b1’; embryos secrete antimicrobial compounds ‘e1-e4’(Fig. 5F) that checks the bacterial growth rate. The amount of embryonic secretion (e1-e4) depends on the LDD in the embryo (Fig. 4A), as a result we find a reciprocal relation between the bacterial growth rate (in ES-BS) and LDD of the embryos from which the ES-BS has been obtained (Fig. 4A). The secretion of e1-e4 by embryos is dependent on the lipolysis of the embryonic LDs (Fig. 4, 5).

We propose that both the embryos and the microbes perceive the presence of one another through the respective secretions of the other. The amount of antimicrobial compound secreted by the embryos depends on amount of LDs. To win the battle over the other, both the microbes and the embryos progressively alter their secretions to counter each other (Middle coulmn in Fig. 4E and last coulmn in Fig. 5D). This mechanism of embryonic defenses is specially functional during its pre-immune stages (0-9 hpf). Fig. 6 is the schematic representation of the mechanism of how the embryos survive in the microbe-laced environment of their aquatic habitat.

**Figure 6.**
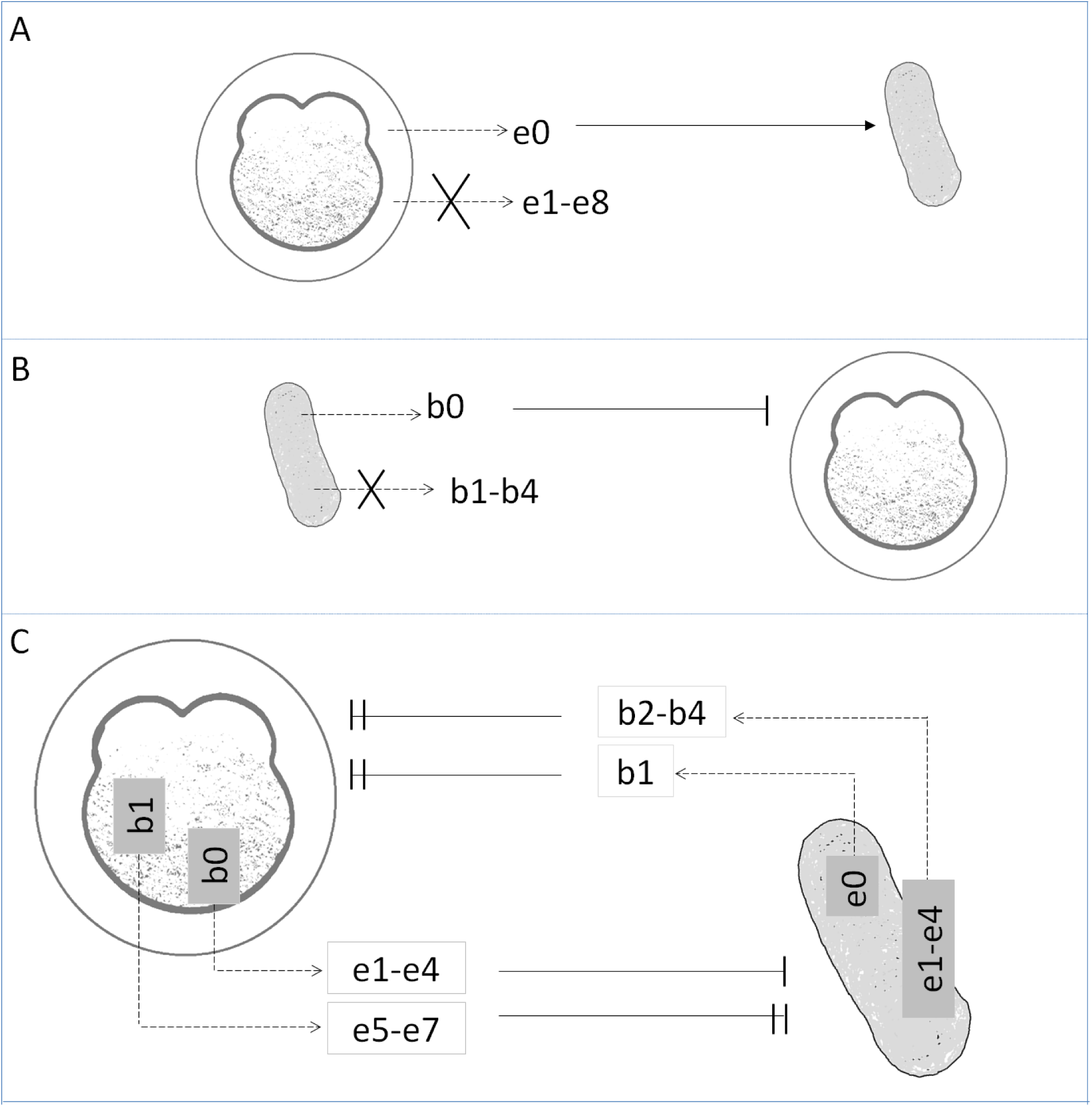
schamatic of the chemical warefare between the preimmune embryo and the microbes. (A) secretions (arrow with dotted line) from the embryos (e0) in microbe free medium has the ability to accelerate the growth rate of the microbes (indicated by the arrow). (B) secretions (dotted arrow) from the microbes (b0) in embryo free medium is detrimental to the development of the embryos (indicated by ‘ **|**’). (C) Response of the embryo and the microbes to microbial (b0-b4) and embryonic (e0-e8) secretions repectively. The microbe secrets b1 (more potent toxin than b0) in response to e0. In response to b0, the embryos secrete antimicrobial compound (e1-e4) however in response to b1 the embryo secretes additional and more potent antimicrobial compound (e5-e7). As a result the microbes secrete even more potent toxins b2-b4 in response to e1-e4.

### Statistical Analyses

All statistical analyses have been performed using the 'pair-sample t-test' in OriginPro 8. The non-significant difference between compared data sets have been designated 'n.s' and signify p>0.05. The significant differences between the data sets fall under three categories: '*' depict p≤0.05, '**' is for p≤0.01 and '***' denote difference between data sets at p≤0.001.

### Experimental procedure

#### 1. Zebrafish handling and embryo culture

Wild type adult zebrafish, *Danio rerio*, was purchased from the local suppliers. The zebrafish were maintained at a temperature of 28°C and an alternate 12 hour light and 12 hour dark cycle. They were housed in a continuous water circualtion system (Aquatic Habitats Inc.). All the experiments were done in agreement with the guide lines provided by the Indian Assocition for the Cultivation fo Science, Animal Ethics Committee. Optimum care was taken at each step to reduce any pain or discomfort to the fish. For breeding, the male and female fish were tranferred to a breeding tank at a ratio of 2:1, respectively, and kept with separators between them overnight. Next day, the separators were removed and the fish were allowed to breed for around 20-30 min. The freshly laid embryos were collected in sterile embryo medium, E3 (50 mM NaCl, 0.17 mM KCl, 0.33 mM CaCl_2_ and 0.33 mM MgSO_4_). All the embryos were maintained in a 28°C incubator until used for further experimentation^26^.

#### 2. Microscopic imaging of the embryos

##### 2.2 Determination of LDD

‘Lipid-Droplet-Density’ (LDD) of the embryos was determined as described in Dutta et.al., 2015^21^. In brief, the embryos were embedded in 0.8% low melting agar (Puregene HiRes Agarose). The embryos were positioned with the animal pole (blastodisc) on top (top view orientation). Time-lapse images were taken at every 10 minutes interval to capture the abundance of LDs and the change in their number over time with embryonic development. Images were acquired using an upright microscope, Olympus BX61, using a water immersion 40X objective. Image processing was done using ImageJ 1.47t software. To determine the embryonic LDD values, we enhanced the contrast of each image first to make the LDs distinctly visible. We then counted the number of LDs visible at each time interval in the embryonic blastodisc. This was divided by the area of the blastodisc visible in each image and expressed as Lipid Droplet Density (LDD) per mm^2^ of blastodisc area. The initial value of LDD was taken as the LDD at 30 mpf (LDD_30_ per mm^2^) to include the time required to prepare the sample and mounting it before imaging.

To determine the ‘rate-of-increase’ of LDD, the LDD values at each time point was plotted against the time points of acquiring the images (corresponding to the developmental time of the embryos) using OriginPRo 8 SR0 software. The slope of each of the plots was evaluated and plotted against the corresponding initial LDD value (LDD_30_) for each embryo. This was done for each case and the average of at least five embryos from each clutch was plotted along with the Standard Error of Mean (S.E.M.) as the error bars.

##### 2.2 Staining and imaging the LDs

To stain the LDs, the embryos were fixed with 4% paraformaldehyde (Merck) followed by repeated wash with Phosphate Buffer Saline (PBS). The LDs were stained using the LD-specific dye, LD540^27^. Post this, LD540 was diluted to 0.1g/ml and the fixed embryos were soaked in it for nearly 20 minutes. The embryos were then washed two to three times with PBS followed by thourough wash with water to remove excess dye which may otherwise cause unnecessary background fluorescence. The embryos were then mounted in 0.6% low melting agar and imaged under 10X magnification using an inverted fluorescence microscope (Zeiss Axio Observer.Z1). The images were processed using the ‘3D-deconvolve’ plugin of ImageJ 1.47t to minimize any unwanted background fluorescence and obtain a 3D-rendered image.

##### 2.3 Determining any physical contact between embryos and bacteria

To determine whether the bacteria come in physical contact with the embryos/LDs, freshly laid embryos (1 cell stage) were agar-embedded and incubated with RFP-labeled bacteria (10^6^/ml *E.coli* DH5α expressing RFP (results are with RFP-bcateria) in E3 media. The embryos were oriented in lateral orientation (side view). Images were acquired by the fluorescence microscope, Zeiss Axio Observer. Z1, under a 10 X objective and green excitation light. Consecutive DIC (for viewing embryo) and fluorescence (for viewing RFP-*E.coli*) images were acquired over time along with z-stacks at each time point. Time-lapse and z-stack images (5 µm step size) of the embryos were acquired at every 20 minutes interval. The z-stack images at each time point were converted to ‘maximum intensity’ projections using ‘3D-projection’ plugin of ImageJ. The images were then deconvolved using ‘3D deconvolve’ plugin of ImageJ. The 3D-rendered overlaid DIC and fluorescence images at each time point give a view of the localization of the bacteria and the embryo as menitoned in Fig. 4A.

### 3 Estimation of embryonic survival and average remaining embryonic survival expectancy

For comparing the survival likelihood of embryos with significantly different LDD_30_ values, freshly laid embryos were collected from different embryo clutches. The survival of the embryos under different treatment conditions was assessed every morning for the entire time of observation (15 dpf in most experiments).

#### 3.1 Survival in the presence or absence of food supply

For comparing the survival in the absence of food for embryos with different LDD_30_ values, embryos were incubated in E3 throughout the entire duration. For the ones to be kept with food, the larvae were fed with larval food from 5 dpf (since zebrafish embryos are lecithotrophic upto 5 dpf). The survival percentage of embryos was assessed at an interval of every 24 hours.

#### 3.2 Survival in the presence or absence of bacteria in medium

To assess embryo survival in the presence or absence of bacterial exposure, freshly laid embryos were collected from different clutches (different LDD_30_ values) and grown in the presence or absence of bacteria in the medium. The survival percentage of embryos was assessed at an interval of every 24 hours.

#### 3.3 Estimation of average remaining embryonic survival expectancy

The average remaining survival expectancy of the embryos denotes the likelihood of survival of the embryos under different treatment conditions. It is calculated using following formula:

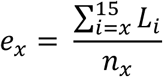

where, *x* is the day of assessment,
*n_x_* is the number of embryos alive at day *x*,

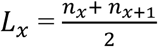

All *e_x_* values mentioned in the manuscript denote the average remaining survival expectancy of the embryos at day zero computed over the data upto fifteen days under different experimentation conditions.

### 4 Determining dry weight of secretion released by the embryos/bacteria

#### 4.1 Embryonic secretions

Embryos were collected at 1-cell stage in E3 medium. For yielding ES-E3 (i.e. embryonic secretions in E3, Table 1) 30 embryos were transferred to 2 ml of fresh E3 and allowed to grow overnight at 28°C. Next morning the embryos were removed and the remaining solution was filtered using a 0.22 μm syringe filter (Merck Millipore). This filtrate was transferred into a pre-weighed microfuge tube. The solution was dried in a rotary-vaccuum evaporator and the tube was re-weighed. The resultant weight obtained upon subtraction of the weight of the pre-weighed empty tube from the weight of the tube with the dried filtrate gives the dry weight of secretion by 30 embryos. This was used to determine weight of secretion released by 1 embryo. For obtaining ES-BS, bacteria was added to E3 and allowed to grow overnight at 37°C. The solution was filtered using syringe filter (this is BS-E3) and this was used to treat 1-cell stage embryos. The rest of the procudre is same as described for ES-E3. For ES-BS(OS), OS was added to BS-E3 and this was used to treat the embryos. Lastly, for ES-BS_ES_, bacteria was grown in previously collected ES-E3 and this filtrate is termed BS_ES_. This was used to treat the embryos. Secretions released were collected in similar manner as described earlier. All experiments involving determination of dry weight of secretions released by the embryos were done using 30 embryos and 2 ml of treatment solution. This were done in triplicates and the mean plotted with the S.E.M. as the error bars.

#### 4.2 Bacterial secretions

Dry weight of bacterial secretions was determined in the same procedure as the embryonic secretions, except that the approximate number of bacteria yielding the dry weight was determined separately and this was used to evaluate dry weight of secretion released per bacterium. For this, the first step is to approximate the number of bacteria in 1 ml culture. We used RFP-*E.Coli* and counted the number of fluorescence dots (per bacteria) under a 10X objective and green excitation light. This value was used to estimate the number of bacteria in 1 ml culture. The slope of the bacterial growth curves (Fig. 6) was multiplied with the respective Optical Density (O.D.) at each time point and this was used to construct a plot of number of bacteria versus time. The Boltzmann growth curve equation which was then fit into the curves is as follows:

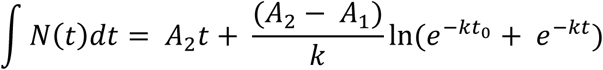

where ʃ*N*(*t*)*dt* is the number of bacteria at time t. t = 24 hours as we evaluate bacterial secretion after 24 hours of incubation under different conditions. Upon fitting and extrapolation we deduce the different parameters. Using this we determine the secretion dry weight from each bacteria. For all experiments involving the treatment of embryos with bacterial secretions, the solution containing the secretions was evaporated in a vacuum drier and the dry weight of the secretion was evaluated. This was re-suspended in E3 to maintain a concentration of 1 mg/ml for all the treatment conditions.

### 5 Drug treatment of the embryos

#### 5.1 Inhibition of lipolysis by Orlistat

The broad spectrum lipase inhibitor, Orlistat (OS) (O-STAT 120, Aristo), was used to inhibit lipolysis in the embryos. As per our previous report, the working concentration of OS i.e. 150 μM is detrimental for the development of the zebrafish embryos and causes death by 6 hpf^25^. Hence, we determined the appropriate OS concentration at which OS treatment does not kill the embryos but delays their overall development (lipolysis still active but at a much lower rate). For this, the embryos were treated with increasing concentrations of OS (from 0 to 150 μM) and the survival of the embryos was assessed after every twenty-four hours of treatment. It is observed that at 10 μM concentration of OS, the embryos survive and hence this was chosen for further experimentation so that, lipolysis is partially inhibited in these embryos, however, they do not die due to the OS treatment.

#### 5.2 Inhibition of protein degradation by Heclin

Heclin (Tocris Bioscience) is a HECT E3 Ub ligase inhibitor^28^. Heclin at its working concentration (7 μM) is detrimental for zebrafish embryos and kills them at around 32-cell stage^25^. Therefore, we determined Heclin concentration at which the embryo survive but the degradation of protein is also inhibited partially. The embryos were treated with varying concentrations of Heclin (0 to 7 μM) and the survival of the embryos was assessed post 24 hours of incubation and the Total Protein Content (TPC) of the embryos was determined for each set. It is found that Heclin at a concentration of 1 μM is not lethal for the embryos but prevents the degradation of protein partially as compared to the control embryos (in E3). Newly laid embryos were then treated with 1 μM Heclin in the presence or absence of BS-E3 and the dry weight of secretion by the embryos was measured at the end of 24 hours of incubation.

#### 5.3 Inhibition of protein translation by Cycloheximide (CHX)

Cycloheximide (Sigma) is a protein translation inhibitor. At its working concentration of 7 μM it is lethal to the embryos^25^. We varied the concentration of CHX (0 to 1 μM) and determined that at 1μM CHX is no longer lethal to the embryos but does induce inhibition of protein translation. 1-cell stage embryos were treated with 1 μM CHX and kept with/without BS-E3 till the embryos are at 16-cell stage (most secretions released within this time) (Fig. 5C). The dry weight of secretion by the embryos was assessed and compared with the control embryos.

### 6. LD isolation

For LD isolation, 150 embryos were taken at 1-cell stage. The embryos were dechorionated with Pronase (Sigma Aldrich) and were then deyolked using deyolking buffer (55mM NaCl, 1.8 mM KCl, 1.25 mM NaHCO_3_)^29^<sup>29</sup>LD isolation was done according to standard protocol^30^ with a few modifications. Post deyolking, 250 mM of sucrose in T-buffer (300 mM glycine, 120 mM potassium gluconate, 100 mM HEPES, 100 mM taurine, 20 mM NaCl, 2.5 mM MgCl_2_, pH 6.8) was added to the blastodisc fraction followed by homogenization by syringe plunging and pestle based homogenization. This was followed by ultracentrifugation in decreasing sucrose gradient of 70%, 55%, 50% and 15% in T-buffer. LDs were collected from the 15-50% layer and used for further experimentation.

### 7. Disc diffusion assay using LDs, embryonic secretions

An overnight culture of E. coli (DH5α) was spread over agar plates using a sterilized L-spreader. The plate was allowed to dry and then inverted and incubated for around 30 min at 37°C till the bacterial solution had dried completely. The determination of the Anti Microbial (AM) activity of the LDs and the embryonic secretions was done by the disc diffusion assay as mentioned by Anand et. al^22^. In brief, 3 mm filter discs were autoclaved and placed on agar (Luria agar, HiMedia). For cytostatic assay, bacteria (separate plates for gram negative, *E.coli* and gram positive, *B.subtilis* bacteria) was spread on the plate and the desired sample to be tested was pipetted on the discs. This was incubated overnight at 37°C for 24 hours. For cytotoxic assay, bacteria was spread on the plate and allowed to grown overnight at 37°C for 24 hours. Next day, the filter discs were placed and the samples were loaded on them. This was again kept for overnight incubation. The clear area around the discs denotes the zone of inhibition of the sample. For control, 20 μl of T-buffer, 20 μl of 250mM sucrose in T-buffer (for LD control), 20μl of DMSO (for LD-lipid control) and standard antibiotic, Ampicillin at the optimum concentration of 50μg/ml were put on the sterile discs.

## Acknowledgements

The authors were financially supported by Ramanujan fellowship, funding from Department of Science and Technology and Department of Biotechnology, government of India. The work has also been supported by the institutional start up grant.

## Author contributions

A.D. performed all the experiments, analyzed the data and prepared the manuscript. S.B. performed experiment, prepared the manuscript. D.K.S. conceptualized the experiments, data analysis and prepared the manuscript.

## References

1 Trede, N. S. & Zon, L. I. Development of T-cells during fish embryogenesis. Dev. Comp. Immunol. 22, 253–263 (1998).

2 Zapata, A., Diez, B., Cejalvo, T., Gutiérrez-De Frías, C. & Cortés, A. Ontogeny of the immune system of fish. Fish Shellfish Immunol. 20, 126–136 (2006).

3 Magadan, S. Pathogen-Host Interactions: Antigenic Variation v. Somatic Adaptations. Results Probl Cell Differ. 57, 235–264 (2015).

4 Lee, J.-W. et al. RAG-1 and IgM Genes, Markers for Early Development of the Immune System in Olive Flounder, Paralichthys olivaceus. Dev. Reprod. 18, 99–106 (2014).

5 Trede, N. S., Langenau, D. M., Traver, D., Look, A. T. & Zon, L. I. The use of zebrafish to understand immunity. Immunity 20, 367–379 (2004).

6 Lieschke, G. J. & Currie, P. D. Animal models of human disease: Zebrafish swim into view. Nat. Rev. Genet. 8, 353–367 (2007).

7 Novoa, B. & Figueras, A. Zebrafish: Model for the Study of Inflammation and the Innate Immune Response to Infectious Diseases. Curr. Top. Innate Immun. II 946, 253–275 (2012).

8 Herbomel, P., Thisse, B. & Thisse, C. Ontogeny and behaviour of early macrophages in the zebrafish embryo. Development 126, 3735–45 (1999).

9 Le Guyader, D. et al. Origins and unconventional behavior of neutrophils in developing zebrafisLe Guyader, Dorothée, Michael J Redd, Emma Colucci-Guyon, Emi Murayama, Karima Kissa, Valérie Briolat, Elodie Mordelet, Agustin Zapata, Hiroto Shinomiya, and Philippe Herbomel. 2007. Blood 111, 132 LP-141 (2008).

10 Herbomel, P., Thisse, B. & Thisse, C. Zebrafish early macrophages colonize cephalic mesenchyme and developing brain, retina, and epidermis through a M-CSF receptor-dependent invasive process. Dev. Biol. 238, 274–288 (2001).

11 Welte, M. A. Expanding roles for lipid droplets. Curr. Biol. 25, 1–24 (2015).

12 Thiam, A. R., Farese, R. V. & Walther, T. C. The biophysics and cell biology of lipid droplets. Nat. Rev. Mol. Cell Biol. 14, 775–786 (2013).

13 Guo, Y., Cordes, K. R., Farese, R. V. & Walther, T. C. Lipid droplets at a glance. J. Cell Sci. 122, 749–752 (2009).

14 Bailey, A. P. et al. Antioxidant Role for Lipid Droplets in a Stem Cell Niche of Drosophila. Cell 163, 340–353 (2015).

15 Cermelli, S., Guo, Y., Gross, S. P. & Welte, M. A. The Lipid-Droplet Proteome Reveals that Droplets Are a Protein-Storage Depot. Curr. Biol. 16, 1783–1795 (2006).

16 Gao, Q. & Goodman, J. M. The lipid droplet–a well-connected organelle. Front. Cell Dev. Biol. 3, 1–12 (2015).

17 den Brok, M. H., Raaijmakers, T. K., Collado-Camps, E. & Adema, G. J. Lipid Droplets as Immune Modulators in Myeloid Cells. Trends Immunol. 39, 380–392 (2018).

18 Cubillos-Ruiz, J. R. et al. ER Stress Sensor XBP1 Controls Anti-tumor Immunity by Disrupting Dendritic Cell Homeostasis. Cell 161, 1527–1538 (2015).

19 Ibrahim, J. et al. Dendritic cell populations with different concentrations of lipid regulate tolerance and immunity in mouse and human liver. Gastroenterology 143, 1061–1072 (2012).

20 Wang, S., Wang, Y., Ma, J., Ding, Y. & Zhang, S. Phosvitin plays a critical role in the immunity of zebrafish embryos via acting as a pattern recognition receptor and an antimicrobial effector. J. Biol. Chem. 286, 22653–22664 (2011).

21 Dutta, A. & Kumar Sinha, D. Turnover of the actomyosin complex in zebrafish embryos directs geometric remodelling and the recruitment of lipid droplets. Sci. Rep. 5, 1–14 (2015).

22 Anand, P. et al. A novel role for lipid droplets in the organismal antibacterial response. Elife 2012, 1–18 (2012).

23 Heck, A. M., Yanovski, J. A. & Calis, K. A. Orlistat, a new lipase inhibitor for the management of obesity. Pharmacotherapy 20, 270–279 (2000).

24 Sternby, B., Hartmann, D., Borgström, B. & Nilsson, Å. Degree of in vivo inhibition of human gastric and pancreatic lipases by Orlistat (Tetrahydrolipstatin, THL) in the stomach and small intestine. Clin. Nutr. 21, 395–402 (2002).

25 Dutta, A. & Sinha, D. K. Zebrafish lipid droplets regulate embryonic ATP homeostasis to power early development. Open Biol. 7, (2017).

26 Avdesh, A. et al. Regular Care and Maintenance of a Zebrafish (Danio rerio) Laboratory: An Introduction. J. Vis. Exp. (2012). doi:10.3791/4196

27 Spandl, J., White, D. J., Peychl, J. & Thiele, C. Live cell multicolor imaging of lipid droplets with a new dye, LD540. Traffic 10, 1579–1584 (2009).

28 Mund, T., Lewis, M. J., Maslen, S. & Pelham, H. R. Peptide and small molecule inhibitors of HECT-type ubiquitin ligases. Proc. Natl. Acad. Sci. 111, 16736–16741 (2014).

29 Link, V., Shevchenko, A. & Heisenberg, C.-P. Proteomics of early zebrafish embryos. BMC Dev. Biol. 6, 1 (2006).

30 Ding, Y. et al. Isolating lipid droplets from multiple species. Nat. Protoc. 8, 43–51 (2013).

